# Efficient generative modeling of protein sequences using simple autoregressive models

**DOI:** 10.1101/2021.03.04.433959

**Authors:** Jeanne Trinquier, Guido Uguzzoni, Andrea Pagnani, Francesco Zamponi, Martin Weigt

## Abstract

Generative models emerge as promising candidates for novel sequence-data driven approaches to protein design, and for the extraction of structural and functional information about proteins deeply hidden in rapidly growing sequence databases. Here we propose simple autoregressive models as highly accurate but computationally extremely efficient generative sequence models. We show that they perform similarly to existing approaches based on Boltzmann machines or deep generative models, but at a substantially lower computational cost. Furthermore, the simple structure of our models has distinctive mathematical advantages, which translate into an improved applicability in sequence generation and evaluation. Using these models, we can easily estimate both the model probability of a given sequence, and the size of the functional sequence space related to a specific protein family. In the case of response regulators, we find a huge number of ca. 10^68^ sequences, which nevertheless constitute only the astronomically small fraction 10^-80^ of all amino-acid sequences of the same length. These findings illustrate the potential and the difficulty in exploring sequence space via generative sequence models.

## I. INTRODUCTION

The impressive growth of sequence databases is prompted by increasingly powerful techniques in data-driven modeling, helping to extract the rich information hidden in raw data. In the context of protein sequences, unsupervised learning techniques are of particular interest: only about 0.25% of the more than 200 million amino-acid sequences currently available in the Uniprot database [1] have manual annotations, which can be used for supervised methods.

Unsupervised methods benefit from evolutionary relationships between proteins: while mutations modify amino-acid sequences, selection keeps their biological functions and their three-dimensional structures remarkably conserved. The Pfam protein family database [2], *e.g.,* lists more than 18,000 families of homologous proteins, offering rich datasets of sequence-diversified but functionally conserved proteins.

In this context, *generative statistical models* are rapidly gaining interest. The natural sequence variability across a protein family is captured via a probability *P*(*a*_1_,…, *a_L_*) defined for all amino-acid sequences (*a*_1_,…,*a_L_*). Sampling from *P*(*a*_1_,…,*a_L_*) can be used to generate new, non-natural amino-acid sequences, which in an ideal case should be statistically indistinguishable from the natural sequences. However, the task of learning *P*(*a*_1_,…, *a_L_*) is highly non-trivial: the model has to assign probabilities to all 20^*L*^ possible amino-acid sequences. For typical proteins of lengths *L* = 50-500, this accounts to 10^65^ —10^650^ values, to be learned from the *M* = 10^3^ — 10^6^ sequences contained in most protein families. Selecting adequate generative model architectures is thus of outstanding importance.

The currently best explored generative models for proteins are so-called coevolutionary models [3], such as those constructed by the Direct Coupling Analysis (DCA) [4–6] or related methods [7–9]. They explicitly model the usage of amino acids in single positions (i.e. residue conservation) and correlations between pairs of positions (i.e. residue coevolution). The resulting models are mathematically equivalent to Potts models [10] in statistical physics, or to Boltzmann machines in statistical learning [11]. They have found numerous fascinating applications in protein biology:

1. The *effect of amino-acid mutations* is predicted via the log-ratio log{*P*(mutant)/*P*(wildtype)} between mutant and wildtype probabilities. Strong correlations to mutational effects determined experimentally via deep mutational scanning have been reported [12, 13]. A resulting fascinating application is the data-driven design of mutant libraries for protein optimization [14, 15].
2. *Contacts between residues* in the protein fold are extracted from the strongest epistatic couplings between double mutations, *i.e.* from the “direct couplings” giving the name to DCA [4]. These couplings are essential input features in the wave of deep-learning (DL) methods for protein-structure prediction, which currently revolutionize the field of protein-structure prediction [16–19].
3. The generative implementation bmDCA [6] is able to *generate artificial but functional amino-acid sequences* [20, 21]. The potential in applications to protein design is immense, but still almost unexplored.

However, Potts models or Boltzmann machines are not the only generative-model architectures explored for protein sequences. Latent-variable models like Restricted Boltzmann machines [22] or Hopfield-Potts models [23] learn dimensionally reduced representations of proteins; using sequence motifs, they are able to capture groups of collectively evolving residues [24] better than DCA models. Variational auto-encoders also achieve dimensional reduction, but in the more flexible setting of deep learning. While their use as generative models for non-natural proteins remains still quite limited [25, 26], they provide currently the best mutational-effect predictors [27]. Deep autoregressive models have been recently proposed for mutational-effect prediction [28], but their generative capacities currently remain unexplored. However, from anecdotal evidence in these works, and in agreement with general observations in machine learning, it appears that deep architectures may be more powerful than shallow architectures, provided that very large datasets and computational resources are available [27]. In general, they are therefore limited to quite small proteins of up to about 200-300 amino acids.

Here we propose an astonishingly simple model architecture called arDCA, based on a shallow (one-layer) autoregressive model paired with generalized logistic regression. Such models are computationally very efficient, they can be learned in few minutes, as compared to days for bmDCA and more involved architectures. Nevertheless, we demonstrate that arDCA provides highly accurate generative models, comparable to the state of the art in mutational-effect and residue-contact prediction. Their simple structure makes them more robust in the case of limited data. Furthermore, and this may have important applications in homology detection [29], our autoregressive models are the only generative models we know about, which allow for calculating exact sequence probabilities, and not only non-normalized sequence weights, thereby enabling the comparison of the same sequence in different models for different protein families.

## II. RESULTS

### A. Autoregressive models for protein families

Protein families are given in form of multiple sequence alignments (MSA) 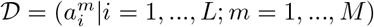 of *M* proteins of aligned length *L*. The entries 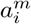 equal either one of the standard 20 amino acids, or the alignment gap In total, we have *q* = 21 possible different symbols in 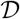. The aim of unsupervised generative modeling is to earn a statistical model *P*(*a*_1_,…,*a_L_*) of (aligned) full-length sequences, which faithfully reflects the variability found in 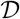: sequences belonging to the protein family of interest should have comparably high probabilities, unrelated sequences very small probabilities.

Here we propose a computationally efficient approach based on *autoregressive models.* We start from the exact decomposition

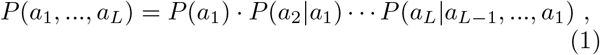

of the joint probability of a full-length sequence into a product of more and more involved conditional probabilities *P*(*a_i_*|*a*_*i*-1_, …,*a*_1_) of single positions *a_i_*, conditioned to all previously seen positions *a*_*i*-1_,…, *a*_1_. While this decomposition is a simple consequence of Bayes’ theorem, it suggests an important change in viewpoint on generative models: while learning the full *P*(*a*_1_, …,*a_L_*) from 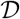 is a task of unsupervised learning (sequences are not labeled), learning the factors *P*(*a*_*i*_|*a*_*i*-1_, …,*a*_1_) becomes a task of supervised learning, with (*a*_*i*-1_,…, *a*_1_) being the input (feature) vector, and *a_i_* the output label (in our case a categorical q-state label). We can thus build on the full power of supervised learning, which is methodologically more explored than unsupervised learning [30–32].

In this work, we choose the following parameterization:

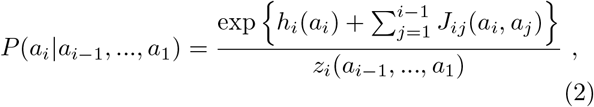

with 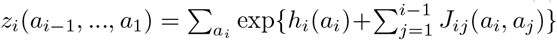 being a normalization factor. In machine learning, this parameterization is known as soft-max regression, the generalization of logistic regression to multi-class labels [31]. This choice, as detailed in the SI, enables a particularly efficient parameter learning by likelihood maximization. Because the resulting model is parameterized by a set of “fields” *h_i_*(*a*) and “couplings” *J_ij_* (*a,b*) as in DCA, we dub our method as “arDCA”.

A few remarks are needed:

- Eq. (2) has striking similarities to standard DCA [5], but also important differences. The two have exactly the same number of parameters, but their meaning is quite different. While DCA has symmetric couplings *J_ij_*(*a,b*) = *J_ji_*(*b,a*), the parameters in Eq. (2) are directed and describe the influence of site *i* on site *j* for *j > i* only, *i.e.* only one triangular part of the matrix is filled.
- The inference in arDCA is very similar to plmDCA [33], *i.e.* to DCA based on pseudolikelihood maximization [34]. In plmDCA each *a_i_* is, however, conditioned to *all* other **a*j* in the sequence, and not only by partial sequences. The resulting directed couplings are usually symmetrized akin to standard Potts models. On the contrary, the *J_ij_*(*a,b*) that appear in arDCA cannot be interpreted as “direct couplings” in the DCA sense, cf. below for details on arDCA-based contact prediction. However, plmDCA has limited capacities as a generative model, because parameter symmetrization causes an accuracy loss. No such symmetrization is needed for arDCA.
- arDCA, contrary to all other DCA methods, allows for calculating the probabilities of single sequences. In bmDCA, we can only determine sequence weights, but the normalizing factor, *i.e.* the partition function, remains inaccessible for exact calculations; expensive thermodynamic integration via MCMC sampling is needed to estimate it. The conditional probabilities in arDCA are individually normalized; instead of summing over *q^L^* sequences we need to sum *L*-times over the *q* states of individual amino acids. This may turn out as a major advantage when the same sequence in different models shall be compared, as in homology detection and protein family assignment [35, 36].

### B. The positional order matters

Eq. (1) is valid for any order of the positions, *i.e.* for any permutation of the natural positional order in the amino-acid sequences. This is no longer true, when we parameterize the *P*(*a_i_*|*a*_*i*-1_, …,*a*_1_) according to Eq. (2). Different orders may give different results. In the SI we show that the likelihood depends on the order, and that we can optimize over orders. We also find that the best orders are correlated to the entropic order, where we select first the least entropic, *i.e.* most conserved, variables, and then the individually most variable positions of highest entropy. The site entropy *s_i_* = - Σ_*a*_ *f_i_*(*a*)log *f_i_* (*a*) can be directly calculated from the empirical amino-acid frequencies *f_i_*(*a*) of all amino acids *a* in site *i*.

Because the optimization over the possible L! site orderings is very time consuming, we use the entropic order as a practical heuristic choice. In all our tests, described in the next sections, the entropic order does not perform significantly worse than the best optimized order we found.

A close-to-entropic order is also attractive from the point of view of interpretation. The most conserved sites come first. If the amino acid on those sites is the most frequent one, basically no information is transmitted further. If, however, a sub-optimal amino acid is found in a conserved position, this has to be compensated by other mutations, *i.e.* necessarily by more variable (more entropic) positions. Also the fact that variable positions come last, and are modeled as depending on all other amino acids, is well interpretable: these positions, even if highly variable, are not necessarily unconstrained, but they can be used to finely tune the sequence to any sub- optimal choices done in earlier positions.

For this reason, all coming tests are done using increasing entropic order, *i.e.* with sites ordered before model learning by increasing empirical *s_i_* values. The SI shows a comparison with alternative orderings, such as the direct one (from 1 to *L*), a random one, and the optimized one, cf. also Table I for some results.

**TABLE I.**
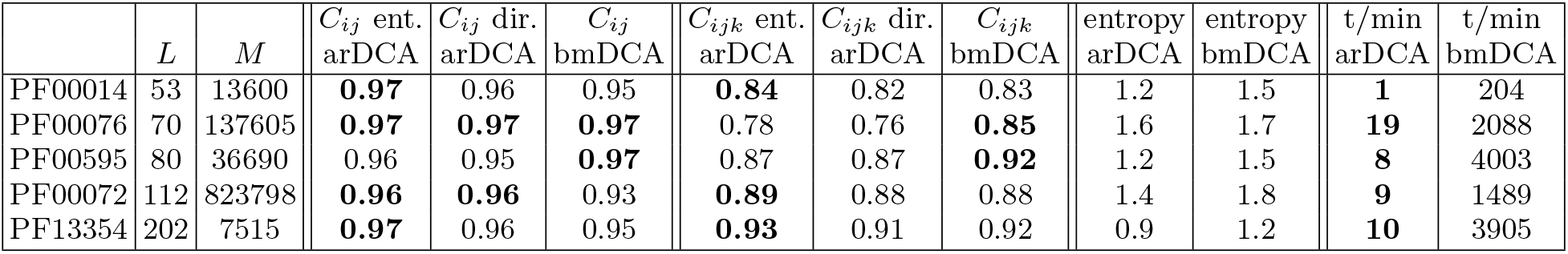
The table summarizes the data used (protein families, sequence lengths *L* and numbers *M*, together with the Pearson correlations between empirical and model-generated connected correlations *C_ij_* and *C_ijk_*, for bmDCA and arDCA using an entropic or direct ordering of sites. The entropies/site and computational running times for model learning are also provided for arDCA and bmDCA. Best values for each measure are evidenced. Similar results for the 32 protein families with deep-mutational scanning data are given in the SI.

### C. arDCA provides accurate generative models

To check the generative property of arDCA, we compare it with bmDCA [6], *i.e.* the most accurate generative version of DCA obtained via Boltzmann machine learning. bmDCA was previously shown to be generative not only in a statistical sense, but also in a biological one: sequences generated by bmDCA were shown to be statistically indistinguishable from natural ones, and most importantly, functional *in vivo* for the case of chorismate mutase enzymes [21].

Here, we compare the statistical properties of natural sequences with those of independently and identically distributed (i.i.d.) samples drawn from *P*(*a*_1_, …,*a_L_*). At this point, another important advantage of arDCA comes into play: while generating i.i.d. samples from a Potts model requires MCMC simulations, which in some cases may have very long decorrelation times and thus become computationally expensive, drawing a sequence from the arDCA model *P*(*a*_1_,…,*a_L_*) is very simple. The factorized expression Eq. (1) allows for sampling amino acids position by position, following the chosen positional order.

Figs. 2A-C show the comparison of the one-point amino-acid frequencies *f_i_*(*a*), and the connected two- and three-point correlations

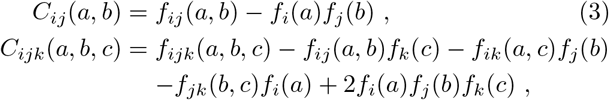

of the data with those estimated from a sample of the arDCA model. Results are shown for the Pfam family PF00076 [2]. Other proteins are shown in Table I and the SI. We find that, for these observables, the empirical and model averages coincide very well, equally well or even slightly better than for the bmDCA case. In particular for the one- and two-point quantities this is quite surprising: while bmDCA fits them explicitly, *i.e.* any deviation is due to imperfect fitting of the model, arDCA does not fit them explicitly, and nevertheless obtains higher precision.

**FIG. 1.**
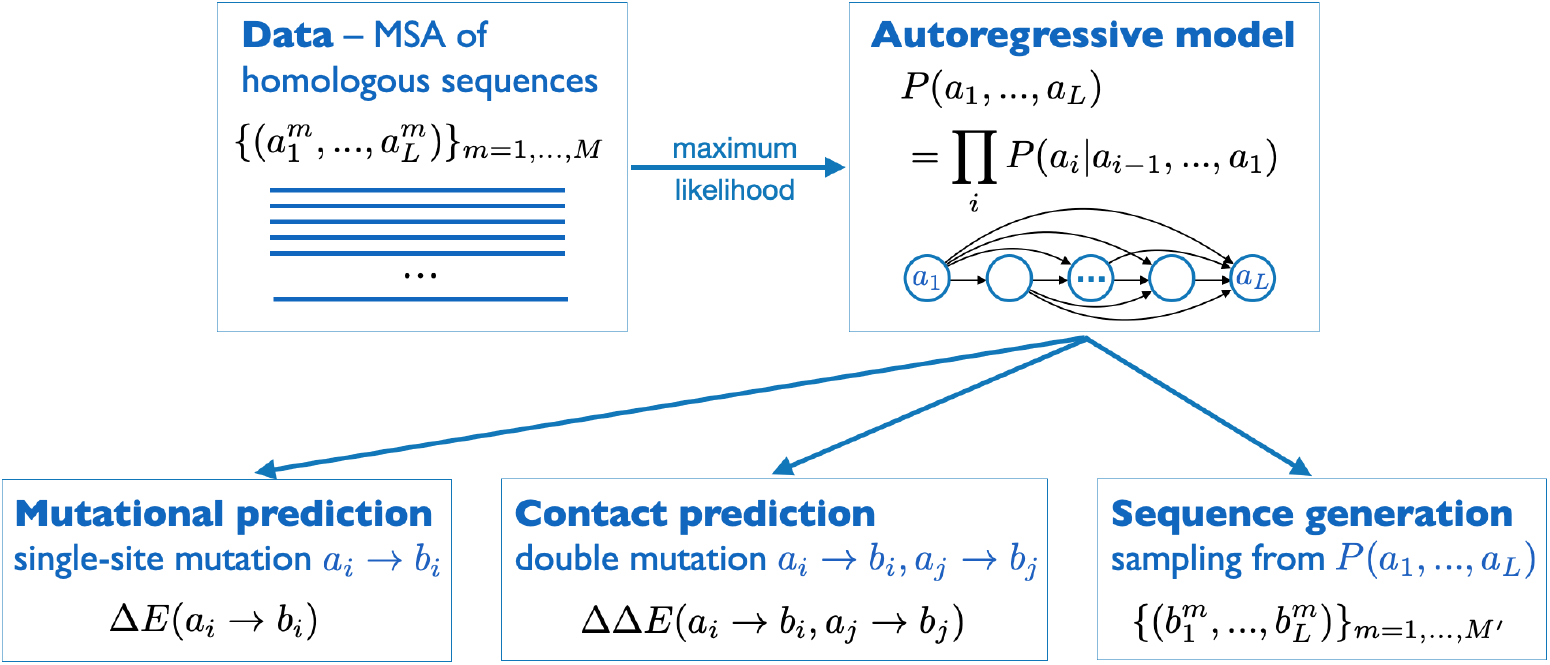
Schematic representation of the arDCA approach: Starting from a MSA of homologous sequences, we use maximum-likelihood inference to learn an autoregressive model, which factorizes the joint sequence probability *P*(*a*_1_,…,*a_L_*) into conditional single-residue probabilities *P*(*a_i_*|*a*_*i*-1_,…, *a*_1_). Defining the statistical energy *E*(*a*_1_,…, *a_L_*) = — log *P*(*a*_1_,…, *a_L_*) of a sequence, we consequently predict mutational effects and contacts as statistical energy changes when substituting residues individually or in pairs, and we design new sequences by sampling from *P*(*a*_1_, …,*a_L_*).

**FIG. 2.**
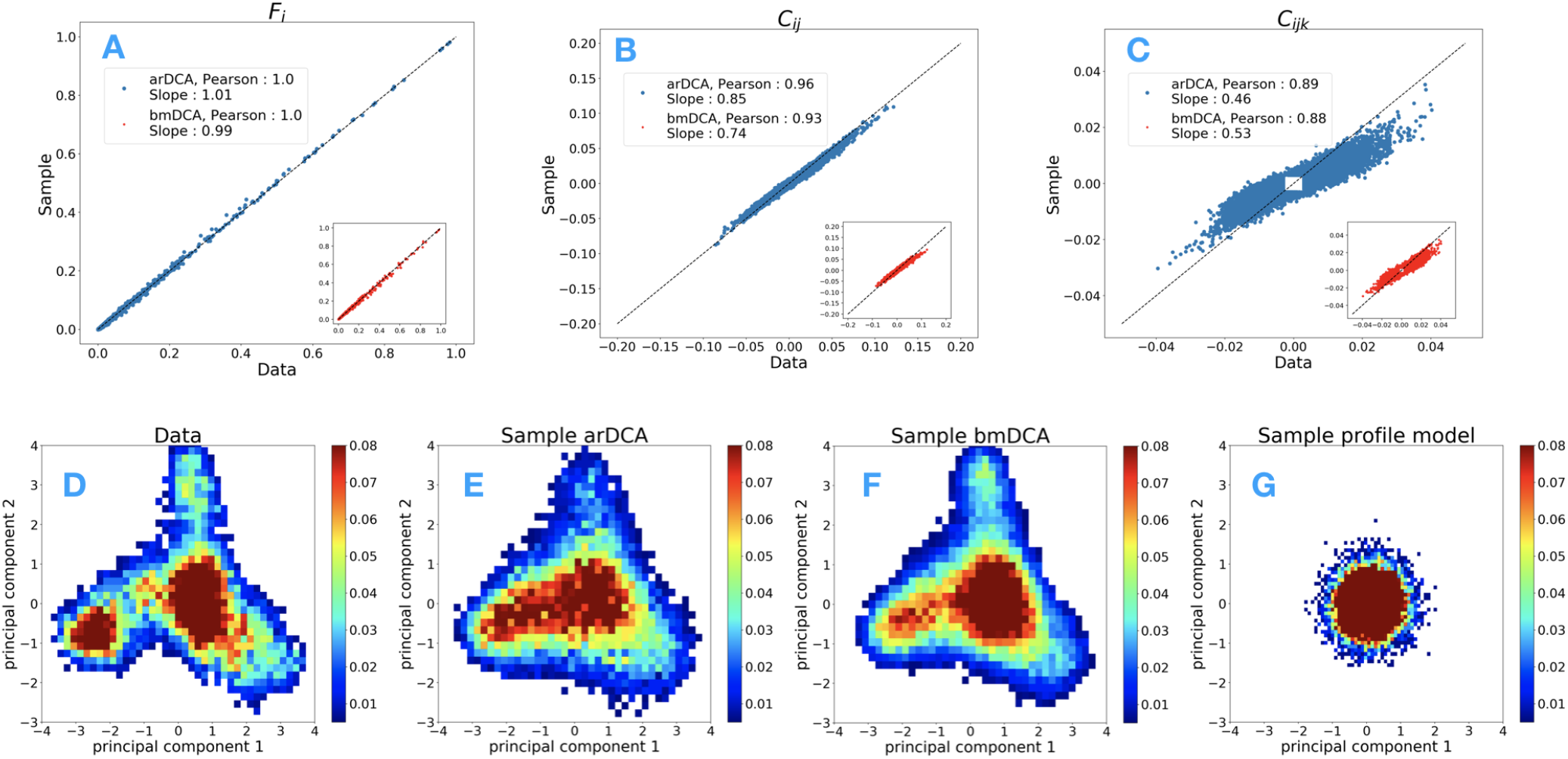
Generative properties of arDCA for PF00076: Panels A-C compare the single-site frequencies *f_i_*(*a*) and two- and three-site connected correlations *C_ij_* (*a,b*) and *C_ijk_* (*a,b,c*) found in the sequence data and samples from models, for arDCA (blue) and bmDCA (red). Panels D-G show different samples projected onto the first two principal components of the natural data. Datasets are the natural MSA (D) and samples from arDCA (E), bmDCA (F) and a profile model (G). Results for other protein families are shown in the SI.

A second test is given by Figs. 2D-G. Panel D shows the natural sequences projected onto their first two principal components (PC). The other three panels show generated data projected onto the same two PCs of the natural data. We see that both arDCA and bmDCA reproduce quite well the clustered structure of the response-regulator sequences (both show a slightly broader distribution than the natural data, probably due to the regularized inference of the statistical models). On the contrary, sequences generated by a profile model *P*_prof_ (*a*_1_,…,*a_L_*) = ∏_*i*_ *f_i_*(*a_i_*) assuming independent sites, do not show any clustered structure: the projections are concentrated around the origin in PC space. This indicates that their variability is almost unrelated to the first two principal components of the natural sequences.

From these observations (cf. also the SI), we conclude that arDCA provides excellent generative models, of at least the same accuracy of bmDCA. This suggests fascinating perspectives in terms of data-guided statistical sequence design: if sequences generated from bmDCA models are functional, also arDCA-sampled sequences should be functional. But this is obtained at much lower computational cost, cf. Table I and without the need to check for convergence of MCMC, which makes the method scalable to much bigger proteins.

### D. Predicting mutational effects via *in-silico* deep mutational scanning

The probability of a sequence is a measure of its “goodness”. For high-dimensional probability distributions, it is generally convenient to work with log-probabilities. Using inspiration from statistical physics, we introduce a “statistical energy”

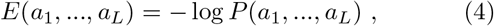

as the negative log-probability. We thus expect functional sequences to have very low statistical energies, while unrelated sequences show high energies. In this sense, statistical energy can be seen as a proxy of (negative) fitness. Note that in the case of arDCA, the statistical energy is not a simple sum over the model parameters as in DCA, but contains also the logarithms of the local partition functions *z_i_*, cf. Eq. (2).

Now, we can easily compare two sequences differing by one or few mutations. For a single mutation *a_i_* → *b_i_*, where amino acid *a_i_* in position *i* is substituted with amino acid *b_i_*, we can determine the statistical-energy difference

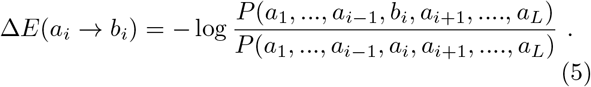

If negative, the mutant sequence has lower statistical energy; the mutation *a_i_* → *b_i_* is thus predicted to be ben-eficial. On the contrary, positive Δ*E* predict a deleterious mutation. Note that, even if not explicitly stated on the left-hand side of Eq. (5), the mutational score Δ*E*(*a_i_* → *b_i_*) depends on the whole sequence background (*a*_1_,…, *a*_*i*-1_, *a*_*i*+1_,…., *a_L_*) it appears in, *i.e.* on all other amino acids *a_j_* in all other positions *j ≠ i*.

It is now easy to perform an *in-silico* deep mutational scan, *i.e.* to determine all mutational scores Δ*E*(*a_i_* → *b_i_*) for all positions *i* = 1, …, *L* and all target amino acids b¿ relative to some reference sequence. In Fig. 3, we compare our predictions with experimental data over more than 30 distinct experiments and wildtype proteins, and with state-of-the art mutational-effect predictors. These contain in particular the predictions using plmDCA (aka evMutation [13]), variational autoencoders (DeepMutation [27]), evolutionary distances between wildtype and the closest homologs showing the considered mutation (GEMME [37]) – all of these methods take, in technically different ways, the context dependence of mutations into account. We also compare it to the context-independent prediction using the above-mentioned profile models.

**FIG. 3.**
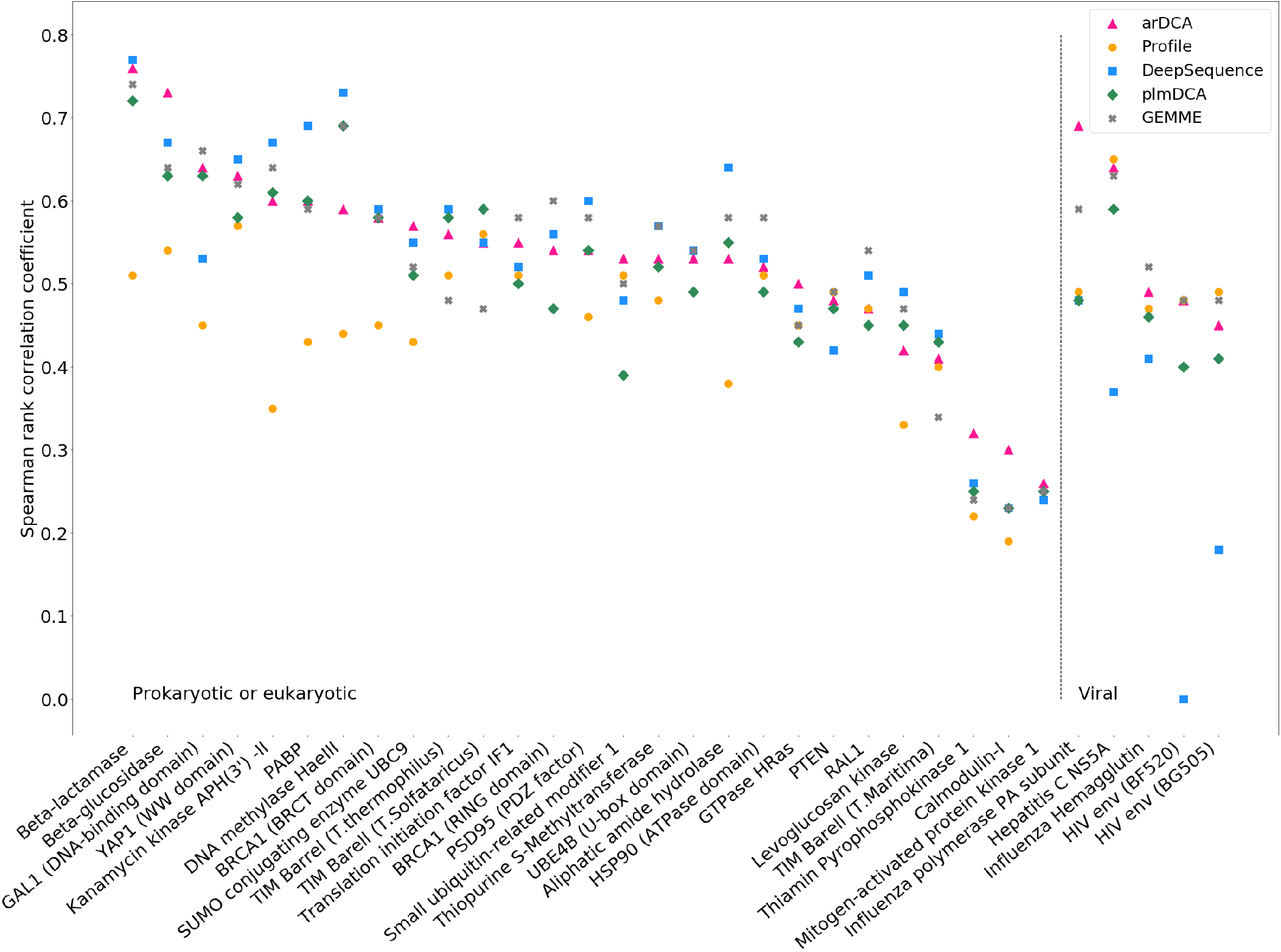
Prediction of mutational effects by arDCA: We show the Spearman rank correlation between results of 32 deep-mutational scanning experiments and various computational predictions. We compare arDCA with profile models, plmDCA (aka evMutation [13]), DeepSequence [27], and GEMME [37], which currently are considered the state of the art. Detailed information about the datasets and the generative properties of arDCA on these datasets are provided in the SI.

It can be seen that the context-dependent predictors outperform systematically the context-independent predictor, in particular for large MSA in prokaryotic and eukaryotic proteins. The four context-dependent models perform in a very similar way; their predictions are usually more correlated to each other than to the experimental data. There is a little but systematic advantage for DeepMutation, which uses the computationally most expensive scheme, and a disadvantage for plmDCA, which was the first published predictor of the ones considered here. GEMME and arDCA perform very similarly.

The situation is a bit different in the typically smaller and less diverged viral protein families. In this case, DeepMutation, which relies on data-intensive deep learning, becomes unstable. It becomes also harder to outperform profile models, *e.g.* plmDCA does not achieve this. arDCA perform similarly or, in one out of four cases, substantially better than the profile model.

We thus find that arDCA permits a very fast and accurate prediction of mutational effect, in line with some of the state-of-the-art predictors. It systematically outperforms profile models and plmDCA, and is more stable than DeepMutation in the case of limited datasets. GEMME, based on phylogenetic informations, astonishingly performs very similarly to arDCA, even if the information taken into account seems different.

As a last remark, we note that for many protein families the Spearman correlation between experimental data and computational predictions is rather limited, with most families in the range of 50-60%. Note that many experiments provide only rough proxies for protein fitness, like *e.g.* protein stability or ligand-binding affinity. To what extent multiple underlying phenotypes can be predicted by unsupervised learning based on homologous MSA thus remains an open question.

### E. Extracting epistatic couplings and predicting residue-residue contacts

The best-known application of DCA is the prediction of residue-residue contacts via the strongest direct couplings [4]. As argued before, the arDCA parameters are not directly interpretable in terms of direct couplings. To predict contacts using arDCA, we need to go back to the biological interpretation of DCA couplings: they represent epistatic couplings between pairs of mutations [38]. For a double mutation *a_i_* → *b_i_,a_j_* → *b_j_*, epistasis is defined by comparing the effect of the double mutation with the sum of the effects of the single mutations, when introduced individually into the wildtype background:

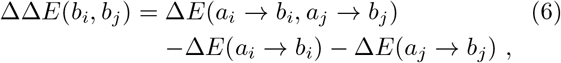

where the Δ*E* in arDCA are defined in analogy to Eq. (5). The epistatic effect ΔΔ*E*(*b_i_, b_j_*) provides an effective direct coupling between amino acids *b_i_, b_j_* in sites *i, j*. In standard DCA, ΔΔ*E*(*b_i_, b_j_*) is actually given by the direct coupling *J_ij_* (*b_i_, b_j_*) — *J_ij_* (*b_i_, a_j_*) — *J_ij_* (*a_i_, b_j_*) + *J_ij_*(*a_i_,a_j_*) between sites *i* and *j*.

For contact prediction, we can treat these effective couplings in the standard way (compute the Frobenius norm in zero-sum gauge, apply the average product correction, cf. the SI for details). The results are represented in Fig. 4. The contact maps predicted by arDCA and bmDCA are very similar, and both capture very well the topological structure of the native contact map. The arDCA method gives in this case a few more false positives, resulting in a slightly lower positive predictive value (panel C). However, note that the majority of the false positives for both predictors are concentrated in the upper right corner of the contact maps, in a region where the largest subfamily of response regulators, the so-called OmpR class, has a homo-dimerization interface.

**FIG. 4.**
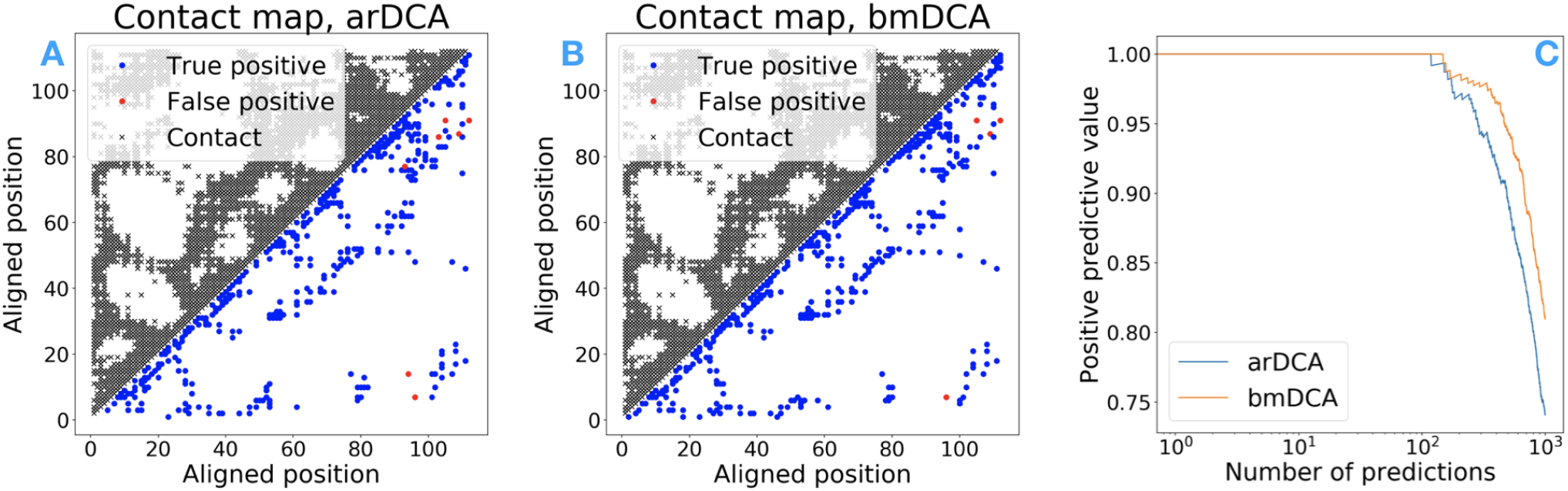
Prediction of residue-residue contacts by arDCA as compared to bmDCA. Panels A and B show the true (black, upper triangle) and predicted (lower triangle) contact maps for PF00076, with blue (red) dots indicating true (false) positive predictions. Panel C shows the positive predictive values (PPV, fraction of true positives in the first predictions) as a function of the number of predictions.

One difference should be noted: for arDCA, the definition of effective couplings via epistatic effects depends on the reference sequence (*a*_1_,…, *a_i_*), in which the mutations are introduced; this is not the case in DCA. So, in principle, each sequence might give a different contact prediction, and accurate contact prediction in arDCA might require a computationally heavy averaging over a large ensemble of background sequences. Fortunately, as is shown in SI, the predicted contacts hardly depend on the reference sequence chosen. It is therefore possible to take any arbitrary reference sequence belonging to the homologous family, and determine epistatic couplings relative to this single sequence. This observation causes an enormous speedup by a factor *M*, with *M* being the depths of the MSA of natural homologs.

### F. Estimating the size of a family’s sequence space

The MSA of natural sequences contains only a tiny fraction of all sequences, which would have the functional properties characterizing a protein family under consideration, *i.e.* which might be found in newly sequenced species or be reached by natural evolution. Estimating this number 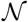 of possible sequences, or their entropy 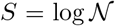, is quite complicated in the context of DCA-type pairwise Potts models. It requires advanced sampling techniques [39, 40].

In arDCA, we can explicitly calculate the sequence probability *P*(*a*_1_, …,*a_i_*). We can therefore estimate the entropy of the corresponding protein family via

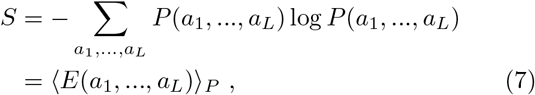

where the second line uses Eq. (4). The ensemble average 〈·〈_*p*_ can be estimated via the empirical average over a large sequence sample drawn from *P*. As discussed before, extracting i.i.d. samples from arDCA is particularly simple due to their particular factorized form.

Results for the protein families studied here are given in Table I. As an example, the entropy density equals *S/L* = 1.4 for PF00072. This corresponds to 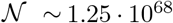 sequences. While being an enormous number, it constitutes only a tiny fraction of all *q^L^* ~ 1.23 · 10^148^ possible sequences of length *L* = 112. Interestingly, the entropies estimated using bmDCA are systematically higher than those of arDCA. On the one hand, this is no surprise: both reproduce accurately the empirical one- and two-residue statistics, but bmDCA is a maximum entropy model, which maximizes the entropy given these statistics [5]. On the other hand, our observation implies that the effective multi-site couplings in *E*(*a*_1_, …,*a_L_*) resulting from the local partition functions *z_i_*(*a*_*i*-1_, …, *a*_1_) lead to a non-trivial entropy reduction.

## III. DISCUSSION

We have presented a class of simple autoregressive models, which provide highly accurate and computationally very efficient generative models for protein-sequence families. While being of comparable or even superior performance to bmDCA across a number of tests including the sequence statistics, the sequence distribution in dimensionally reduced principal-component space, the prediction of mutational effects and residue-residue contacts, arDCA is computationally much more efficient than bmDCA. The particular factorized form of autoregressive models allows for exact likelihood maximization. It allows also for the calculation of exact sequence probabilities (instead of sequence weights for Potts models). This fact is of great potential interest in homology detection using coevolutionary models, which requires to compare probabilities of the same sequence in distinct models corresponding to distinct protein families.

The importance of accurate generative models becomes visible via our results on the size of sequence space (or sequence entropy). For the response regulators used as example throughout the paper (and similar observations are true for all other protein families we analyzed), we find that “only” about 10^68^ out of all possible 10^148^ aminoacid sequences of the desired length are compatible with the arDCA model, and thus suspected to have the same functionality and the same 3D structure of the proteins collected in the Pfam MSA. This means that a random amino-acid sequence has a probability of about 10^-80^ to be actually a valid response-regulator sequence. This number is literally astronomically small, corresponding to the probability of hitting one particular atom when selecting randomly in between all atoms in our universe. The importance of a good coevolutionary modeling becomes even more evident when considering all proteins being compatible with the amino-acid conservation patterns in the MSA: the corresponding profile model still results in an effective sequence number of 10^94^, *i.e.* a factor of 10^26^ larger than the sequence space respecting also coevolutionary constraints. As was verified in experiments, conservation provides insufficient information for generating functional proteins, while taking coevolution into account leads to finite success probabilities.

Finally, autoregressive models can be easily extended by adding hidden layers in the ansatz for the conditional probabilites *P*(*a_i_*|*a*_*i*-1_,…, *a*_1_), with the aim to increase the expressive power of the overall model. For the families explored here, we found that the one-layer model Eq. (2) is already so accurate, that adding more layers only results in similar, but not superior performance, cf. SI. However, in longer or more complicated protein families, the larger expressive power of deeper autoregressive models could be helpful. Ultimately, the generative performance of such extended models should be assessed by testing the functionality of the generated sequences in experiments similar to [21].

## ACKNOWLEDGMENTS

We thank Indaco Biazzo, Matteo Bisardi, Elodie Laine, Anna-Paola Muntoni, Edoardo Sarti and Kai Shimagaki for helpful discussions and assistance with the data. Our work was partially funded by the EU H2020 Research and Innovation Programme MSCA-RISE-2016 under Grant Agreement No. 734439 InferNet. J.T. is supported by a PhD Fellowship of the i-Bio Initiative from the Idex Sorbonne University Alliance.

## Author contributions

A.P., F.Z. and M.W. designed research; J.T., G.U. and A.P. performed research; J.T., G.U., A.P., F.Z. and M.W. analyzed the data; J.T., F.Z. and M.W. wrote the paper.

## Competing interests

The authors declare no competing interests.

## Code availability

Codes in Python and Julia are available at https://github.com/pagnani/ArDCA.git.

## S-I INFERENCE OF THE PARAMETERS

We first describe the inference of the parameters via likelihood maximization. In a Bayesian setting, with uniform prior (we discuss regularization below), the optimal parameters are those that maximize the probability of the data, i.e. a MSA of length *L* with *M* sequences:

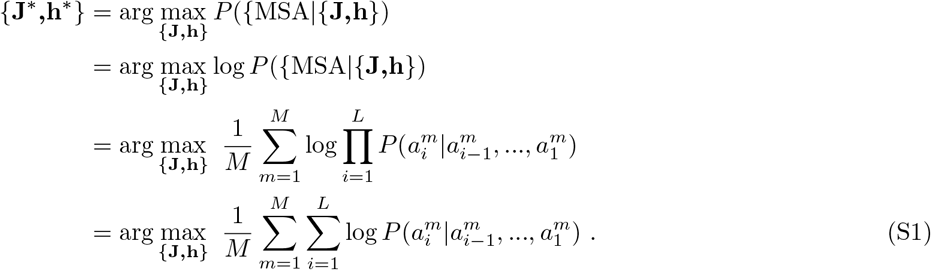

Each parameter *h_i_*(*a*) or *J_ij_*(*a, b*) appears in only one conditional probability *P*(*a_i_*|*a*_*i*-1_,…, *a*_1_), and we can thus maximize independently each conditional probability in Eq. (S1):

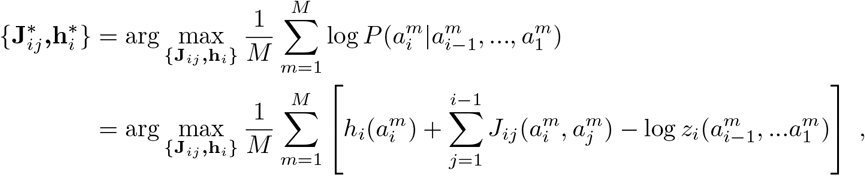

where

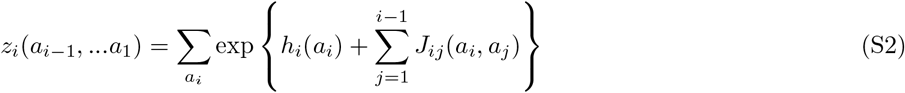

is the normalization factor of the conditional probability of variable *a_i_*.

Differentiating with respect to *h_i_*(*a*), *J_ij_*(*a,b*), we get the set of equations:

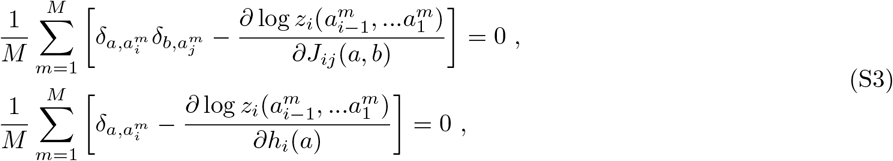

where *δ_a,b_* is the Kronecker delta, and from Eq. (S2):

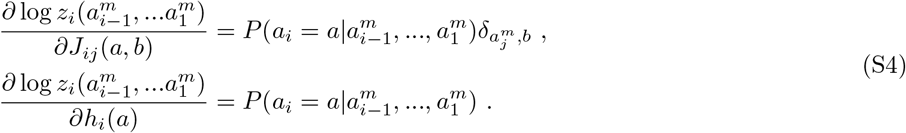

The set of equations thus reduces to a very simple form:

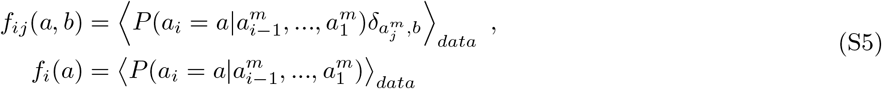

where 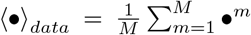 denotes the empirical average, and *f_i_*(*a*), *f_ij_*(*a,b*) are the empirical one- and two-point amino acid frequencies. Note that for the first variable (*i* = 1), which is unconditioned, there is no equation for the couplings, and the equation for the field takes the simple form *f*_1_(*a*) = *P*(*a*_1_ = *a*), which is solved by *h*_1_ (*a*) = log *f*_1_ (*a*) + const.

Unlike the corresponding equations for the Boltzmann learning of a Potts model [6], there is a mix between probabilities and empirical averages in Eq. (S5), and there is no explicit equality between one- and two-point marginals and empirical one and two-point frequencies. This means that the ability to reproduce the empirical one- and two-point frequencies is already a statistical test for the generative properties of the model, and not only for the fitting quality of the current parameter values.

The inference can be done very easily with a simple algorithm using gradient descent, which updates the fields and couplings proportionally to the difference of the two sides of Eq. (S5). We also add a *L*2 regularization, with regularization strength of 0.0001 for the generative tests and 0.001 for mutational effects and contact prediction. Note that the gradients are computed exactly at each iteration, as an average over the data of explicit expressions, hence without the need of MCMC, which provides an important advantage over Boltzmann machine learning.

Finally, in order to partially compensate for the phylogenetic structure of the MSA, which induces correlations among sequences, each sequence is reweighted by a coefficient *w_m_* [5]:

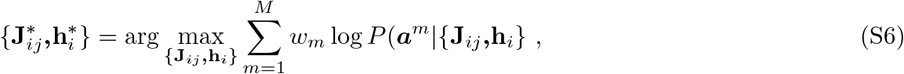

which leads to the same equations as above with the only modification of the empirical average as 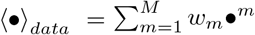. Typically, *w_m_* is given by the inverse of the number of sequences that have at least 80% of residues in common with the sequence m. The goal is to remove the influence of very closely related sequences. Note however that such reweighting cannot fully capture the hierarchical structure of phylogenetic relations between proteins. The effective sequence number *M*_eff_ Σ_*m*_ *w_m_* gives the number of sequences that are actually treated as i.i.d in the inference.

## S-II. SAMPLING FROM THE MODEL

Once the model parameters are inferred, because in this model all the conditional probabilities are known, a sample can be generated by the following procedure:

- Sample the first residue from *P*(*a*_1_)
- Sample the second residue from *P*(*a*_2_|*a*_1_) where *a*_1_ is sampled in the previous step
- … Sample the last residue from *P*(*a_L_*|*a*_*L*-1_,*a*_*L*-2_,…, *a*_2_, *a*_1_)

**FIG. S1.**
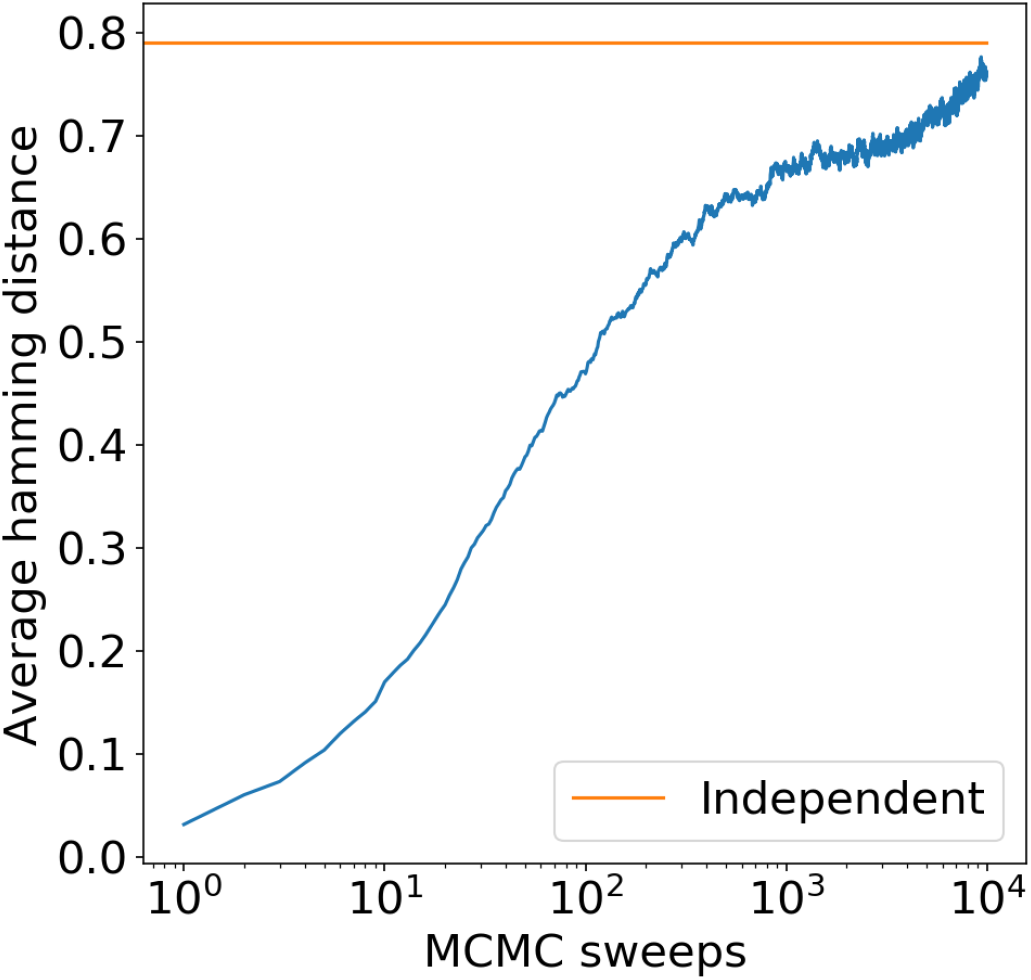
Averaged Hamming distance between a sequence and its time evolution after MCMC sweeps. The average is made with respect to 100 initially equilibrated sequences. The horizontal line is the average Hamming distance between two independent equilibrium sequences.

Each step is very fast because there are only 21 possible values for each probability. On the contrary, sampling from the bmDCA model is more difficult, because a sequences must be obtained via MCMC sampling, but a lot of moves have to be made in order to achieve a proper equilibration in some families [6].

To emphasize the advantages of this direct sampling protocol, we consider the Pfam family PF13354, which is particularly hard to sample via MCMC [41]. In order to directly compare with MCMC sampling of bmDCA, we sampled sequences from our arDCA model, using a Metropolis-Hasting procedure. We propose a random change of a residue, and we accept the move with a probability that depends on the ratio of the probabilities of the new and old sequences. This is of course a very inefficient way of sampling from the arDCA model, but it allows for a direct comparison with the MCMC dynamics of a bmDCA. The Hamming distance between an initial equilibrium configuration (obtained via the sequential procedure described above, which thus guarantees equilibration) and its time evolution after MCMC sweeps was computed. This time-dependent Hamming distance, averaged over 100 initial sequences, is reported in Figure S1. Its shape is very similar to that obtained by MCMC sampling of bmDCA [41]. It grows very slowly with time, and only at very long times it saturates to the equilibrium Hamming distance between two independently sampled sequences. The time it takes to reach this plateau gives an estimation of the number of MCMC sweeps needed to obtain an equilibrium sample. Figure S1 shows that the equilibration takes at least 10^4^ MCMC sweeps. On the other hand, the sequential procedure described above, which is only possible for arDCA models, allows one to sample almost instantaneously, thus completely bypassing the long time scale associated to MCMC. We note that this observation has interesting implication for the problem of sampling in disordered systems with slow dynamics, as already noted in [42].

## S-III. CONTACT PREDICTION

Once the effective couplings are calculated, the standard procedure of DCA is applied [5]. First, each pair of amino acids is assigned an interaction score given by the Frobenius norm:

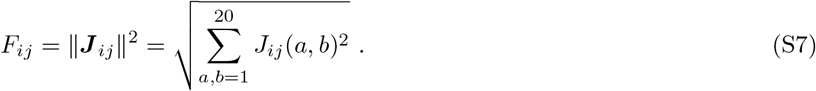

Note that the gap state (*q* = 21) is not taken into account in the norm. Because of overparametrization caused by the non-independence of the empirical frequencies, both the Potts model and autoregressive models are invariant under some gauge transformations, that is different sets of parameters give the same probability. On the other side, the Frobenius norm is not invariant by gauge transformation, so a gauge choice is needed. The zero-sum gauge was found to be the gauge that minimizes the Frobenius norm; in other words, this gauge choice includes in the couplings only the information that cannot be treated by fields. Note that the zero-sum gauge is also the gauge of the standard Ising model. The equations characterizing the zero-sum gauge are: 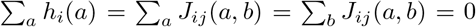. Finally, the so-called average product correction (APC) is applied on the Frobenius scores, because it was empirically shown to improve contact prediction: 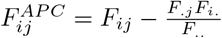 where the dot represents the average with respect to the index.

## S-IV. RESULTS FOR OTHER FAMILIES

### A. Pfam Datasets

#### 1. Description

We describe the properties of the five Pfam families used to test the generative properties and structure prediction of the arDCA model: PF00014, PF00072, PF00076, PF00595, PF13354. The value of *M*_eff_ defined in Section S-I gives the effective number of sequences, obtained by a proper reweighing of very similar sequences.

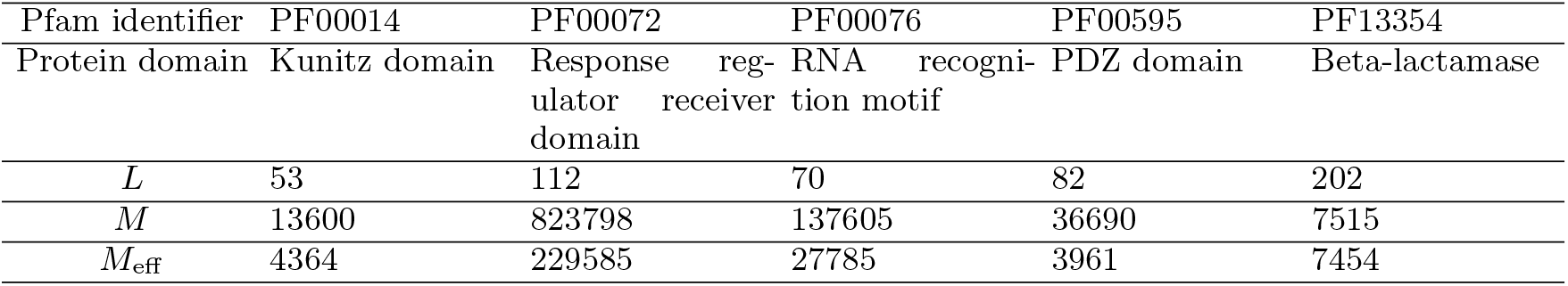

#### 2. Principal component analysis

Figure S2 shows the projection of natural sequences (first column), sequences sampled from arDCA (second column), bmDCA (third column) and the profile model (last column) in a two-dimensional space, constructed by performing principal component analysis on the natural sequences. Each bin in the figure has a color related to its total weight, defined by resampled the sequences using the weights defined in Section S-I.

#### 3. Frequencies

Figure S3 shows how well the model is able to reproduce the empirical frequencies obtained from the data. The one-point frequencies (left), two-point (center) and three-point (right) connected correlations are shown, both from the arDCA model (blue) and the bmDCA (red). Note that for the three-point connected correlations, the correlations that have an empirical value smaller than 0.003 are removed, because they are not meaningful given the limited number of sequences in the dataset.

#### 4. Contact prediction

*a. PPV* To test the contact prediction, a distance *d*(*i,j*) between each heavy atoms in the amino acids was extracted from the crystal structures present in the PDB database. Sites with atoms at a distance < 8 Å were considered in contact. Note that 8 A is too large to be a true contact, but since we are looking for consensus contacts in the family and there is variability from protein to protein, this definition has become standard. Coherently with the literature standard, a minimal separation of |*i — j*| ≥ 5 along the protein chain was imposed in order to consider only non-trivial contacts corresponding to sites that are not close in the chain. Pairs ij are ranked according to the APC-corrected Frobenius norm, as defined in Section S-III. Figure S4 shows the positive predictive value as a function of the number of predicted non-trivial contacts for the arDCA model (blue) and bmDCA (orange). The Positive Predicted Value (PPV) is the fraction of true contacts among the first n predictions, corresponding to the n highest scores.

**FIG. S2.**
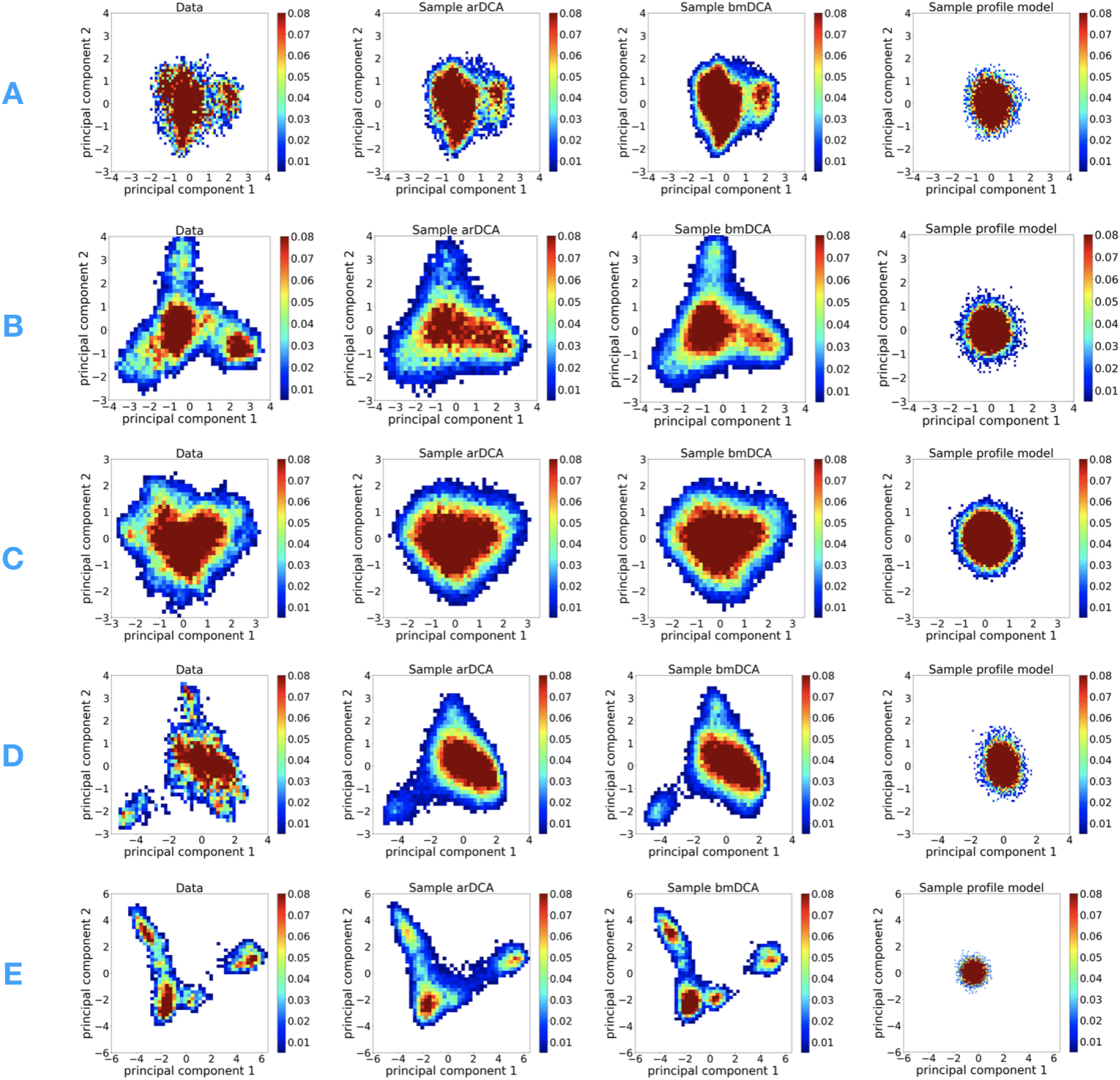
Projections of sequences on the principal components obtained from natural sequences, for the Pfam families PF00014 (A), PF00072 (B), PF00076 (C), PF0595 (D), and PF13354 (E).

*b. Contact map* Figure S5 show the contact maps of the arDCA and bmDCA models. The black crosses represent the true contact map with a threshold of 8 Å. The blue dots are the true positive predictions and the red ones are the false positive considering the 2*L* top predictions of non-trivial contacts.

**FIG. S3.**
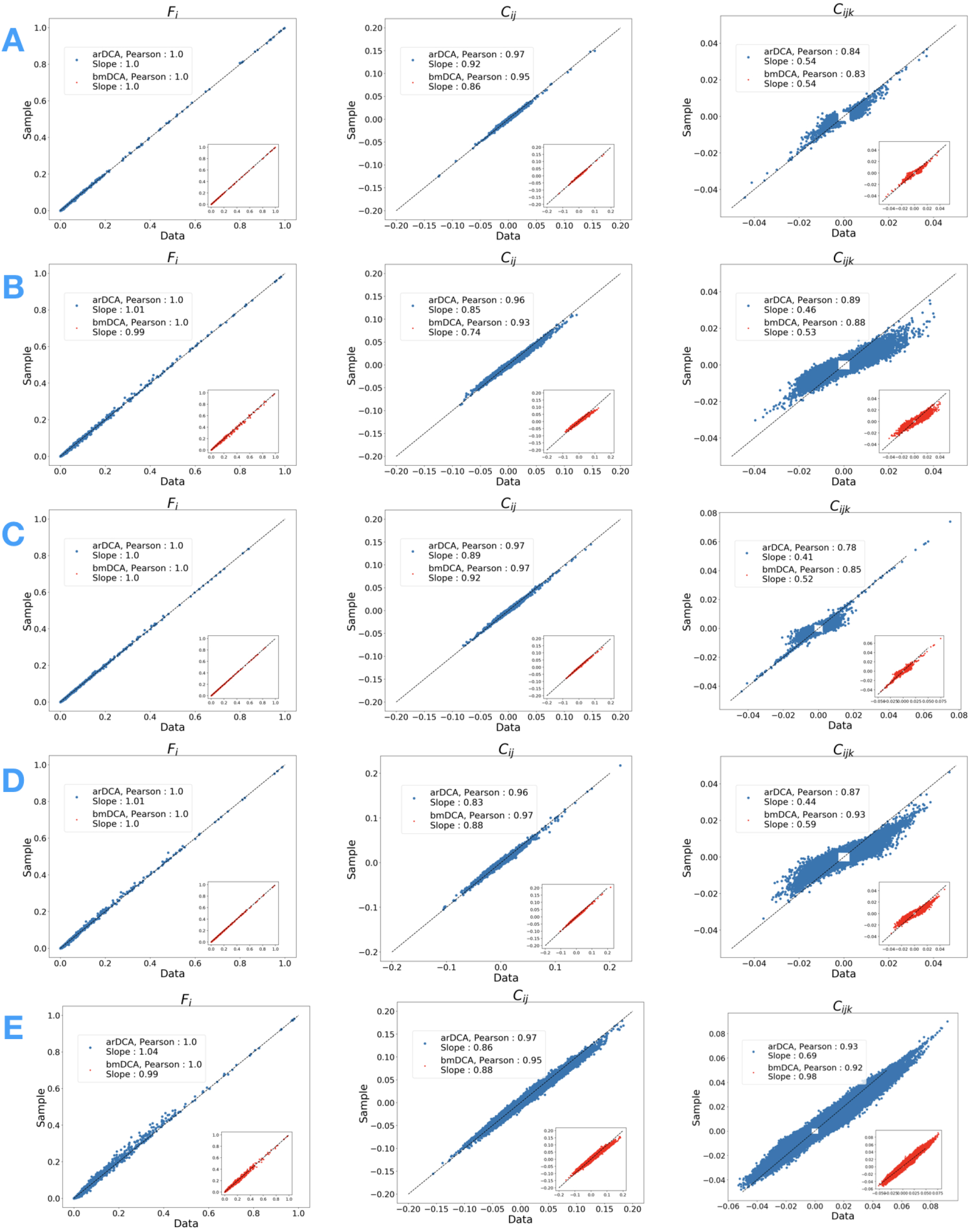
One-point frequencies and two- and three-point connected correlations, obtained from resampling the models (vertical axis) and from empirical data (horizontal axis) for the Pfam families PF00014 (A), PF00072 (B), PF00076 (C), PF0595 (D), and PF13354 (E).

**FIG. S4.**
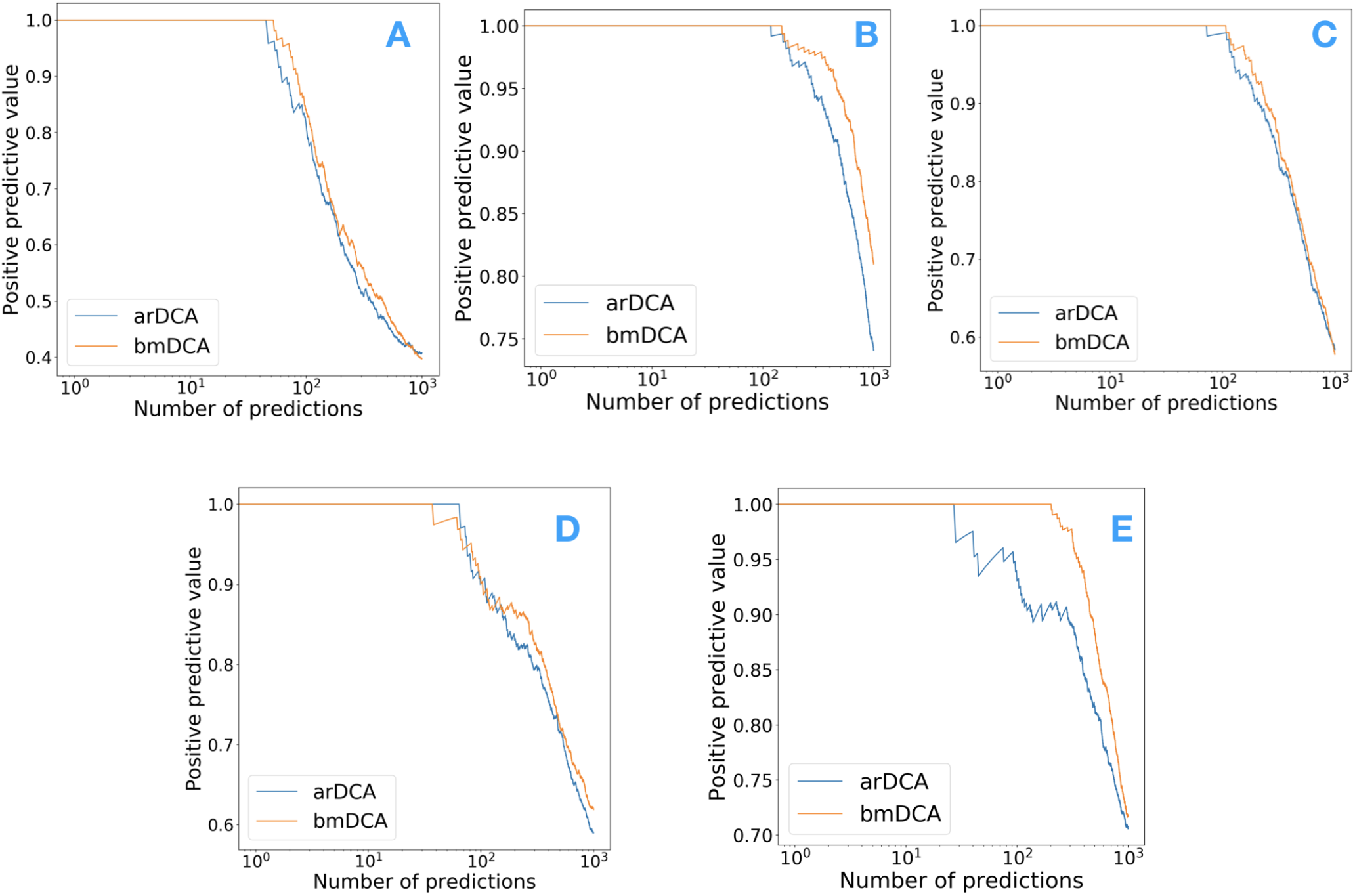
PPV curves for the Pfam families PF00014 (A), PF00072 (B), PF00076 (C), PF0595 (D), and PF13354 (E).

**FIG. S5.**
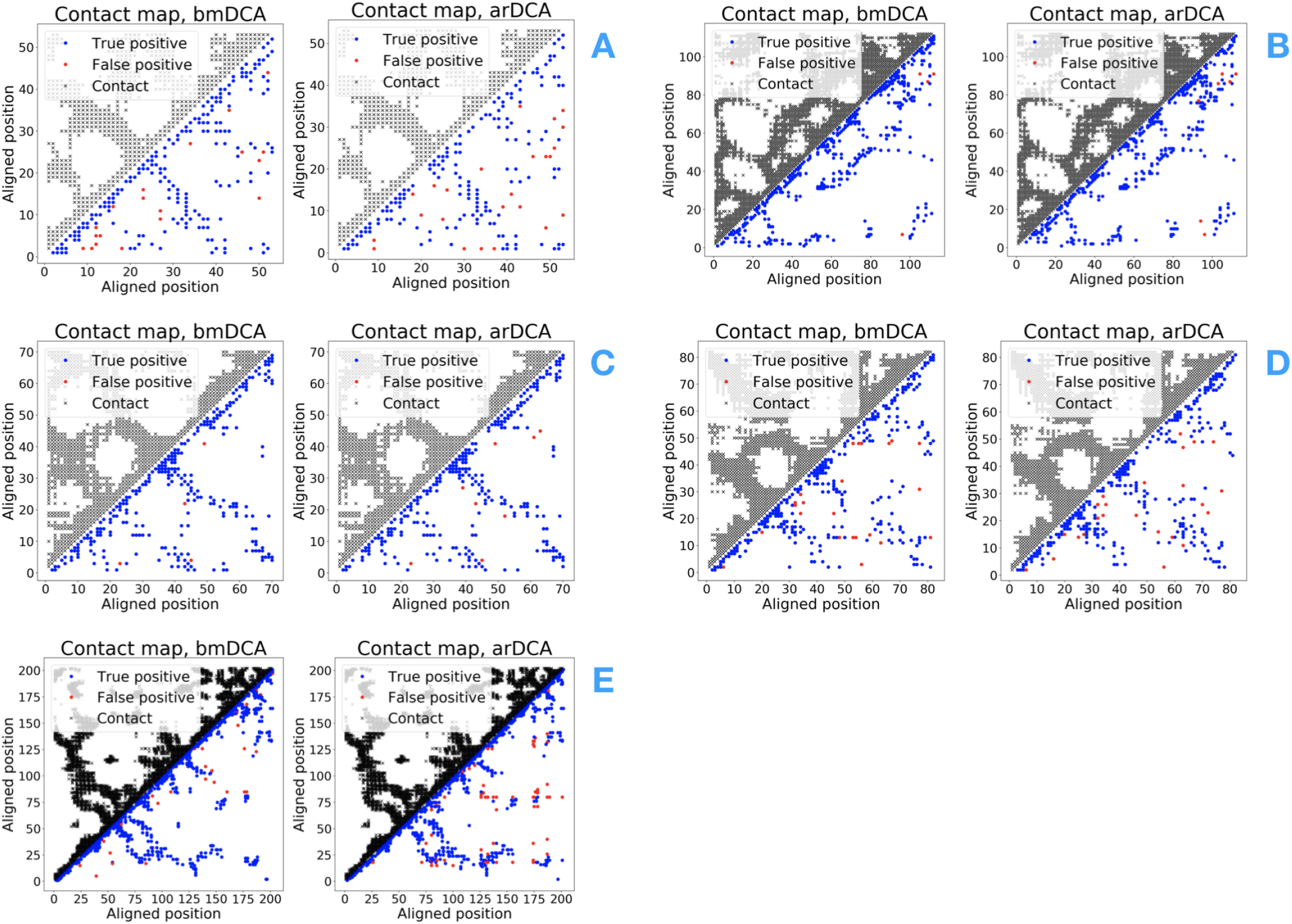
Contact maps for the Pfam families PF00014 (A), PF00072 (B), PF00076 (C), PF0595 (D), and PF13354 (E).

### B. Families used for mutational effects

We show here the generative properties of the arDCA model for the 33 families that are used for mutational effect predictions [27, 37]. The computational time of parameter learning on a standard laptop is also included.

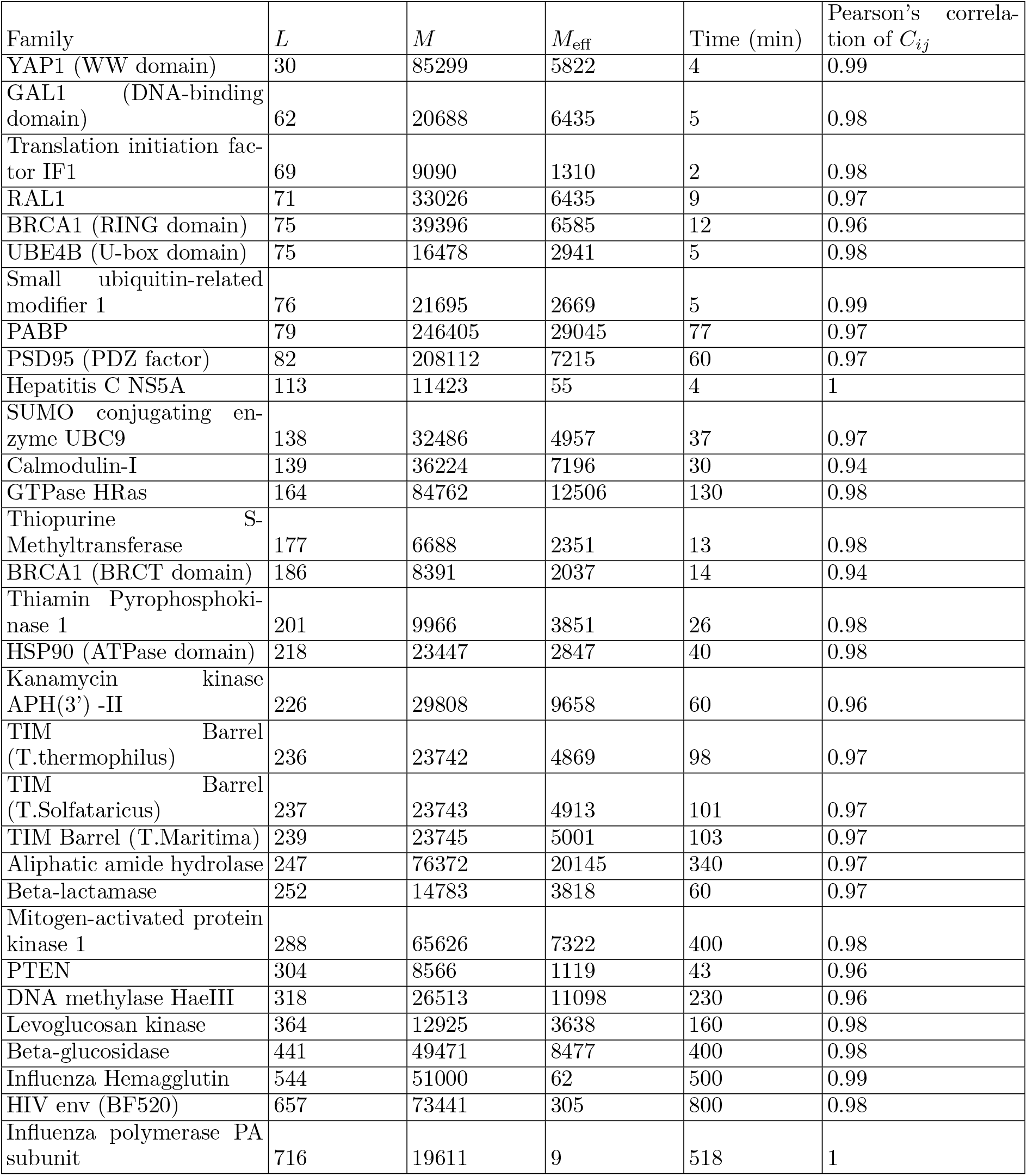

**FIG. S6.**
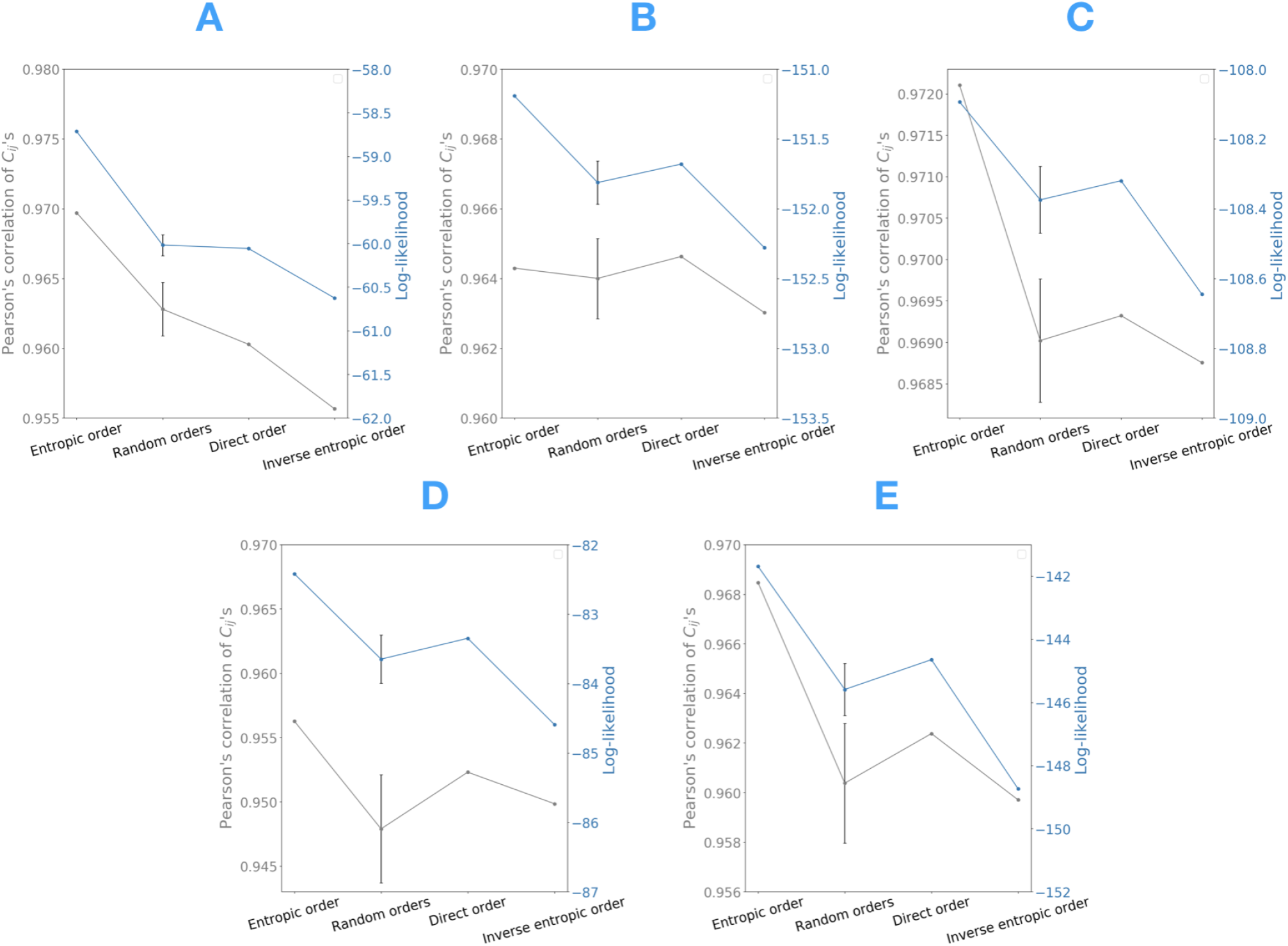
Comparison of the Pearson correlation of two-point connected correlations and the log-likelihood between different orders. The value of the random order is the mean of 100 random orders with one standard deviation given as a vertical bar. Results are for Pfam families PF00014 (A), PF00072 (B), PF00076 (C), PF0595 (D), and PF13354 (E).

## S-V. POSITIONAL ORDER

For a family of length *L*, there are *L*! possible permutations of the sites and therefore *L*! possible orders. The parameterization of the conditional probabilities of the arDCA model is not invariant under a change of order, thus different orders may give different results. However, an optimization over all the different orders is not computationally feasible. We compared some particular orders: the natural order along the protein chain, the entropic order where the sites are ordered in ascending order according to their local entropy 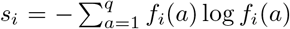, and the inverse entropic order where *s_i_* is used in descending order. A comparison with 100 random orders was also made. The quality of the generative properties was found to be highly correlated with the log-likelihood of the optimized model, which can be computed exactly after the parameters are inferred, see Section S-I. Figure S6 does a comparison of the likelihood and the Pearson’s correlation of the two-point statistics for the different orders. The values reported for the random order is average over the 100 different realizations, with one standard deviation given by the vertical bar. While the direct order is compatible with a random order, for all but one family the entropic order has the highest value of the likelihood and maximizes the Pearson correlation.

In order to check that the entropic order is a good heuristic choice between all possible orders, we designed a greedy procedure to increase the likelihood by doing some permutations between the sites. This procedure tries to find a locally optimal order. The different steps of the procedure are:

- Choose a site randomly
- Permute this site with all the other sites and compute the likelihood of the new model each time
- Choose the permutation that increases the most the likelihood and iterate the procedure

**FIG. S7.**
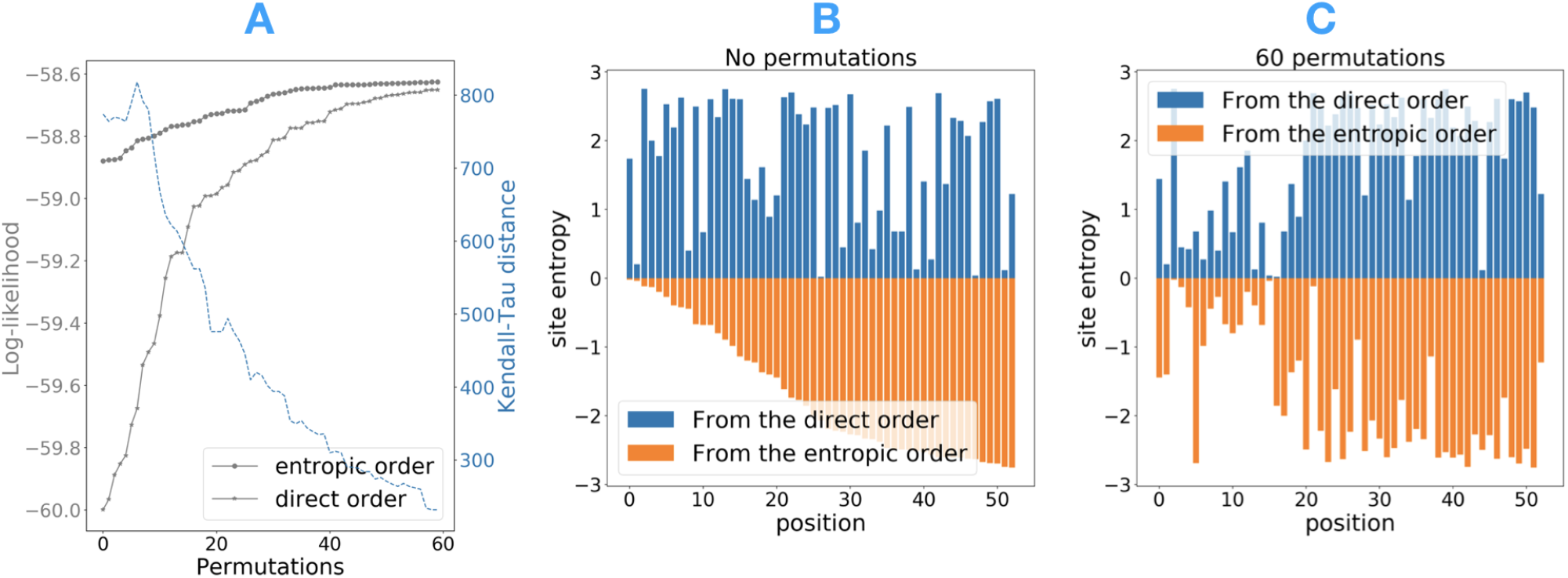
A: Evolution of the log-likelihood and the Kendall-Tau distance of the direct and entropic orders under permutations. B and C: Values of the entropy of each site in the direct and entropic orders with no permutations (B) and after 60 permutations (C).

Figure S7A shows the evolution of the log-likelihood of the entropic and direct orders of the family PF00014 under permutations. The permuted entropic order saturates quickly to a value of the log-likelihood relatively close to the initial one, indicating that the entropic order is not far from a locally optimal one. The direct order saturates to the same value of the log-likelihood. The plot also shows the evolution of the Kendall-Tau distance between the two orders, defined as the number of pairs in a different order, i.e for two lists *l*1 and *l*2,

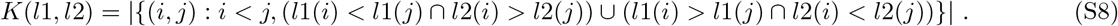

The Kendall-Tau distance gives a measure of the dissimilarity between two lists. Figures S7A shows that the distance between the two orders decreases with increasing permutations. Figures S7B and S7C show the value of the local entropy for each site in both orders before (B) and after (C) 60 permutations. After 60 permutations, it is clear that the sites with a low local entropy are typically at the beginning, which is coherent with the explanation given in the main text.

Finally, Figure S8 shows the Pearson correlation of two-point connected correlations for the 33 families used for mutational effects with the entropic and the direct order. Coherently with the previous discussion, the entropic order gives a better result for 30 over 33 families.

## S-VI. TWO LAYERS AUTOREGRESSIVE MODELS

Due to the very simple structure of the one-layer arDCA model, one might ask whether a more complicated and flexible model could perform better. To address this question, we considered an arDCA model for the family PF00076, but with two layers instead of one. As an exploratory step, we just considered very simple two-layers architectures: each conditional probability *P*(*a_i_*|*a*_*i*-1_,…, *a*_1_), *i* ∈ 2,…, *L* is modeled in terms of a dense input node of size *kq* × (*i* — 1)*q* for (input amino acid sequences are one-hot-encoded) with non a linear activation function *σ*, while the second layer is again a dense node of size *q × kq* concatenated with a *softmax* to get a probability as final output:

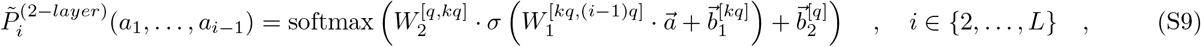

where 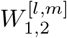 is are parameter matrices of size *l × m* and 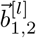 are vectors of parameters (biases) of size *l* that are optimized in the training step. We tried different values of *k*, and different types of activation functions *σ* and we opted for *k* = 5 and *σ* = ReLU.

**FIG. S8.**
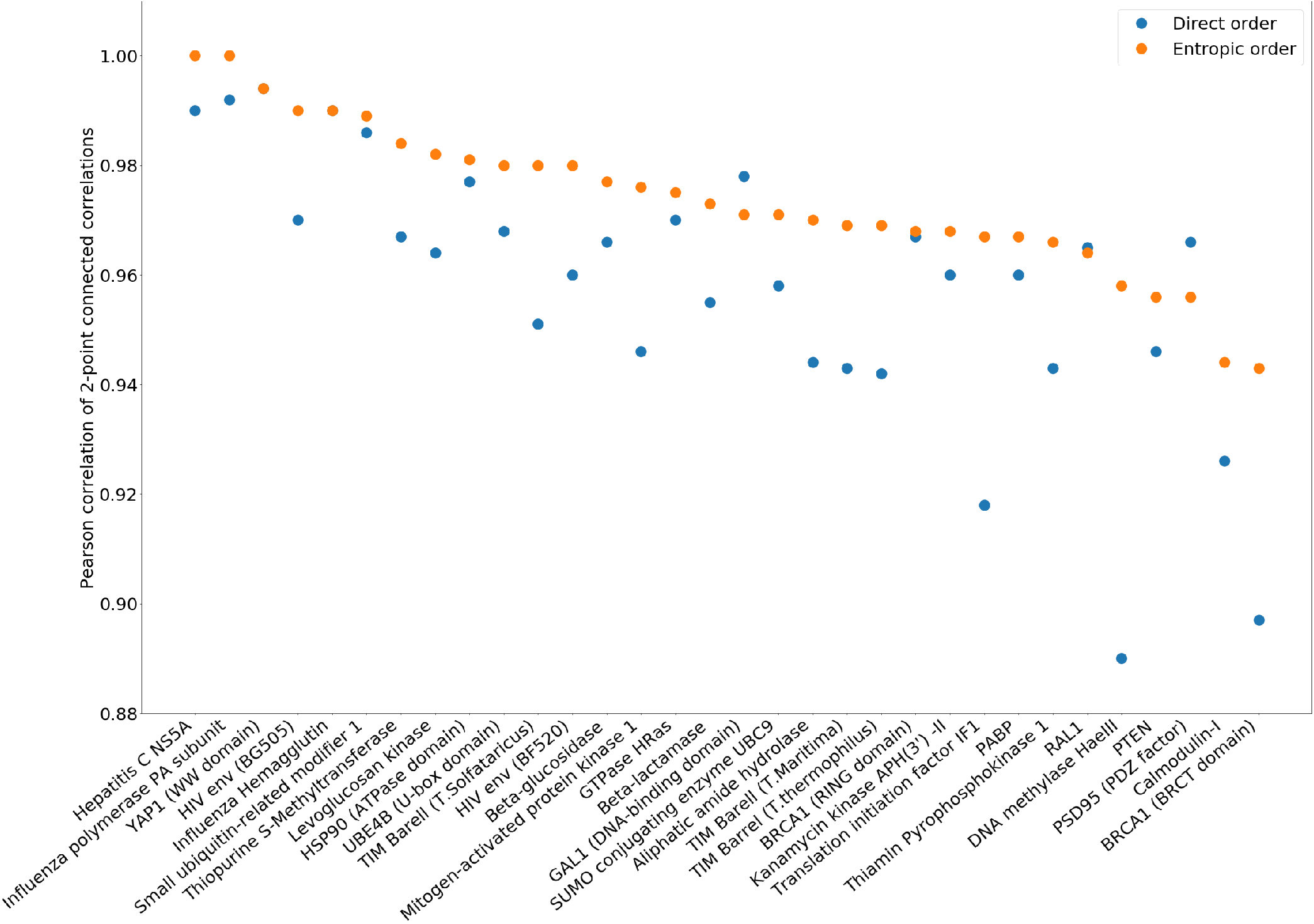
Pearson correlation of the two-point connected correlations for all the 33 families used for mutational effects, for the direct and entropic orders.

Figure S9 shows that increasing the complexity of the model does not improve the ability to reproduce the statistics of the data. In the specific case of the family PF00076, the one-layer model is even marginally better, both for the one-point and two-point statistics. Moreover, the computational time is comparable in the two cases, therefore using a two-layer model gives neither an advantage on the generative qualities, nor on the computational time.

**FIG. S9.**
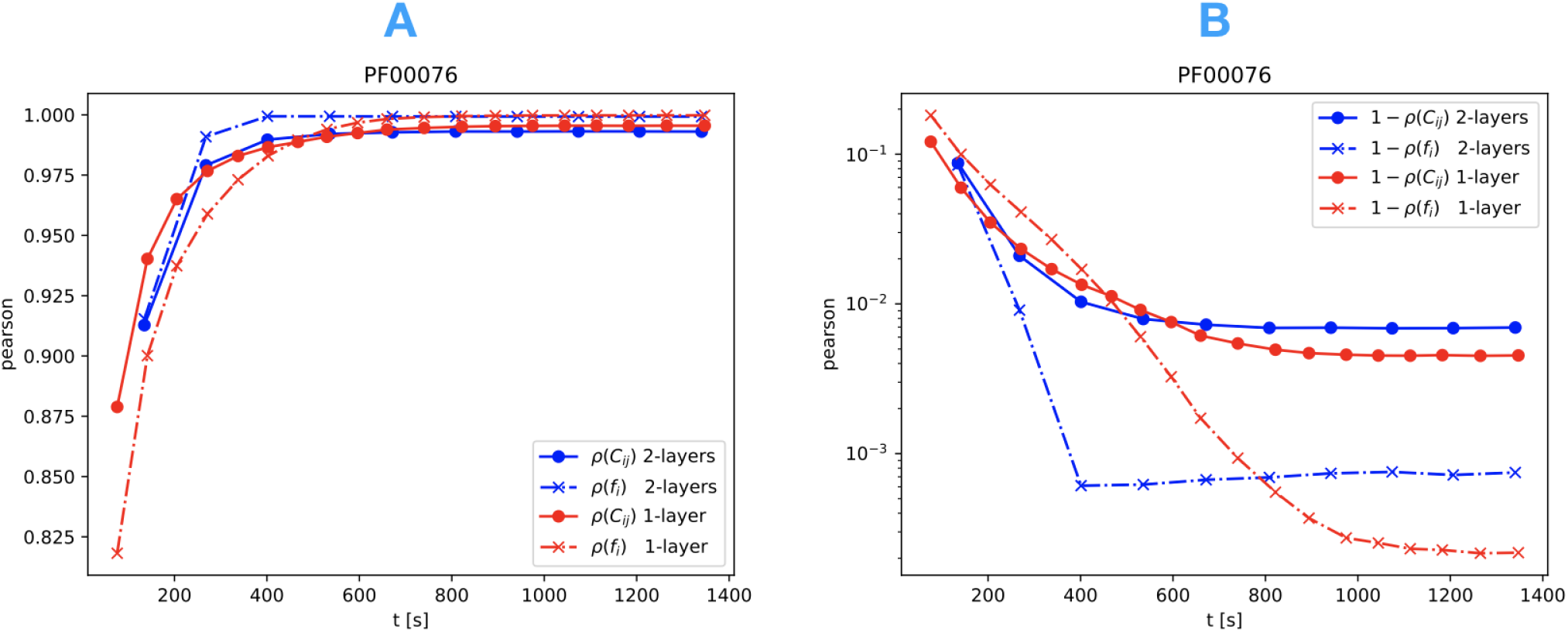
Evolution of the Pearson correlation (for one- and two-point statistics) during the learning for the family PF00076. The computational time is given in seconds. Both figures compare the one-layer (red) and the two-layer (blue) models. Figure A shows the Pearson correlation while figure B shows one minus the Pearson correlation (in semilog scale).

## Notes

### Competing Interest Statement

The authors have declared no competing interest.

